# Topological Data Analysis reveals robust alterations in the whole-brain and frontal lobe functional connectomes in Attention-Deficit/Hyperactivity Disorder

**DOI:** 10.1101/751008

**Authors:** Zeus Gracia-Tabuenca, Juan Carlos Díaz-Patiño, Isaac Arelio, Sarael Alcauter

**Author notes:** Corresponding author: Sarael Alcauter, Instituto de Neurobiologia, Lab C-12, Universidad Nacional Autonoma de Mexico, Blvd Juriquilla 3001, Queretaro, Mexico, 76230.

## Abstract

The functional organization of the brain network (connectome) has been widely studied as a graph; however, methodological issues may affect the results, such as the brain parcellation scheme or the selection of a proper threshold value. Instead of exploring the brain in terms of a static connectivity threshold, this work explores its algebraic topology as a function of the filtration value (i.e., the connectivity threshold), a process termed the Rips filtration in Topological Data Analysis. Specifically, we characterized the transition from all nodes being isolated to being connected into a single component as a function of the filtration value, in a public dataset of children with attention-deficit/hyperactivity disorder (ADHD) and typically developing children. Results were highly congruent when using four different brain segmentations (atlases), and exhibited significant differences for the brain topology of children with ADHD, both at the whole brain network and at the functional sub-network levels, particularly involving the frontal lobe and the default mode network. Therefore, this approach may contribute to identify the neurophysio-pathology of ADHD, reducing the bias of connectomics-related methods.

**Highlights:**

- Topological Data Analysis was implemented in functional connectomes.
- Betti curves were assessed based on the area under the curve, slope and kurtosis.
- The explored variables were robust along four different brain atlases.
- ADHD showed lower areas, suggesting decreased functional segregation.
- Frontal and default mode networks showed the greatest differences between groups.

**Graphical Abstract:** 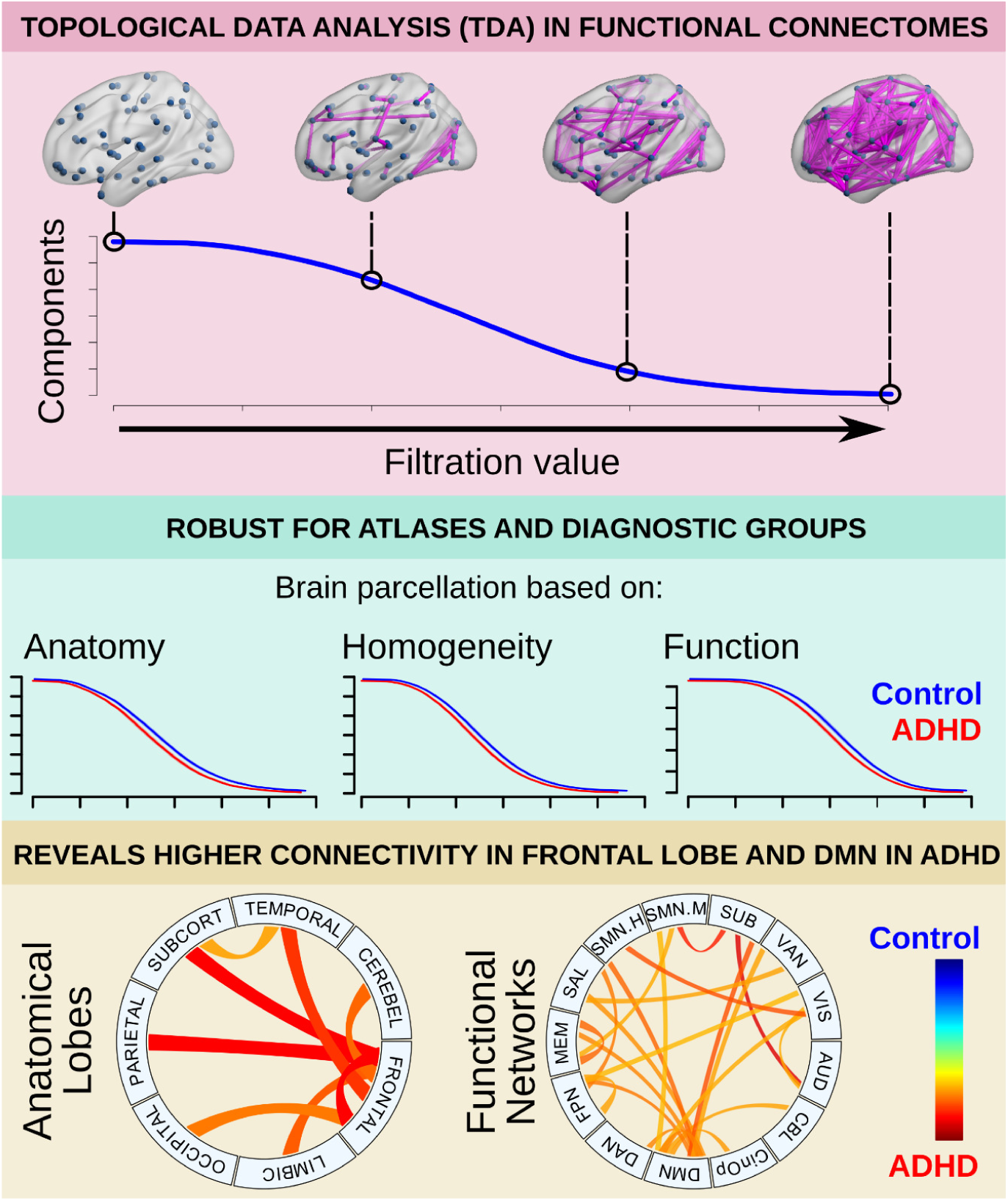

## 1. Introduction

Current neuroimaging technology allows the exploration of the human brain as a network of structurally and/or functionally connected constituents, i.e., voxels or regions of interest. In particular, functional connectivity is defined as the synchrony of neuronal activity patterns of anatomically separated brain regions (Aertsen et al., 1989; Friston et al., 1993) and many studies have explored this property to provide new insights about the functional organization of the brain in health and disease (Van Den Heuvel and Pol, 2010; Lee et al., 2013; Lord et al., 2017), providing the means to study the neurofunctional alterations of neurological and psychiatric disorders from a systems perspective.

One of the most commonly used frameworks to explore the functional brain network is graph theory, which provides a theoretical basis to describe and characterize complex networks (Rubinov and Sporns, 2010; Fornito et al., 2013). In this framework, the brain network is modeled as a graph composed of a set of nodes (mainly voxels or larger regions) and their connections (in this case, the functional connectivity between pairs of elements). In practice, this is constructed using a matrix where each entry is a measure of connectivity between two nodes and then a threshold is applied in order to construct an adjacency matrix which represents the non-spurious connections. However, there is no general criterion to assign an appropriate set of regions of interest (ROIs), nor a defined threshold, which may result in divergent results among studies. For instance, several studies exploring the functional connectome of children diagnosed with Attention-Deficit/Hyperactivity Disorder (ADHD), have reported different results. Specifically, some studies have found higher network segregation and lower integration in ADHD patients compared to controls (Wang et al., 2009; Lin et al., 2014), while others found no differences when exploring the same properties (Cocchi et al., 2012; Sato et al., 2013). Such divergent results may be partially explained by the variability in methods, including threshold and ROIs selection (Konrad and Eickhoff, 2010; Castellanos and Aoki, 2016), as well as the variable robustness of some of the most used approaches (Somandepalli et al., 2015).

Recently, topological data analysis (TDA), has been adopted in neuroimaging as a tool to quantify and visualize the evolution of the brain network at different thresholds (Lee et al., 2011, 2017; Sizemore et al., 2018, 2019; Expert et al., 2019). The main objective of this method is to model the network as a topological space instead of a graph (Edelsbrunner et al., 2000; Zomorodian and Carlsson, 2005), allowing the assessment of the functional connectivity matrix as a topological process instead of a static threshold-dependent representation of the network. One of the possible applications is to characterize how the isolated nodes gradually bind together into larger components (sets of connected nodes) as a function of the filtration value (connectivity threshold), until a single component or simplicial complex is recruited. For this purpose, the number of components at a given filtration value is termed the Betti-0 (see Methods). This process is summarized in a so called Betti-0 curve (Figure 1), which has been shown to differentiate children with developmental disorders from controls using data from positron emission tomography and defining the brain network at the group-level (Lee et al., 2011, 2012). However, the consistency of the method across different brain segmentation schemes has not been explored. Furthermore, it has been typically applied to brain networks defined at the group level, i.e. exploring the covariance of concatenated physiological or structural data from groups of subjects, instead of exploring the individual characteristics of the network and comparing them between groups.

**Figure 1.**
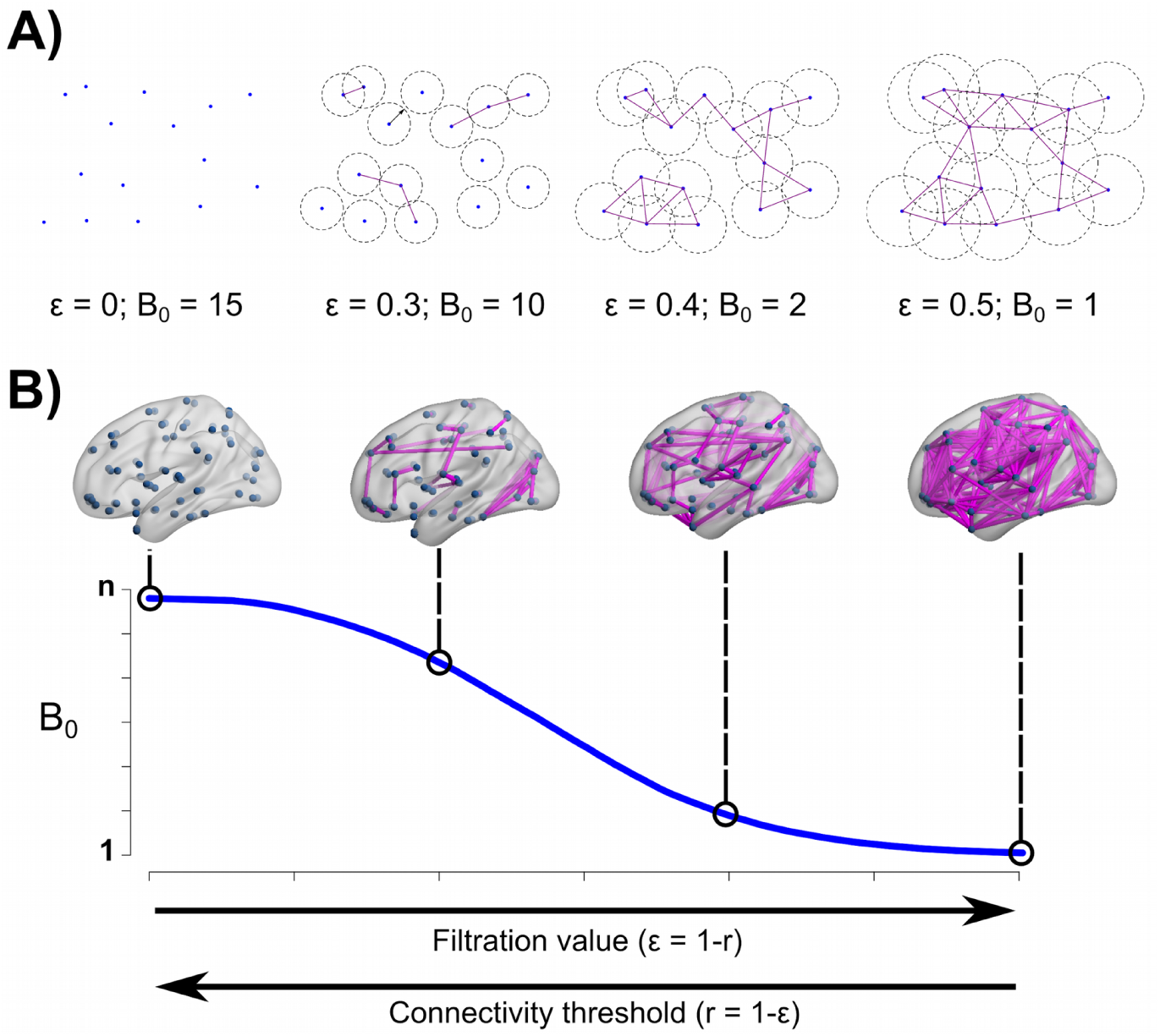
The Betti-0 curve. A) A set of 15 nodes, four filtration values ε, represented as the circle diameter and their corresponding Betti-0 (B_0_). B) Betti-0 curve for a hypothetical brain network; each point in the curve represents the B_0_ for each filtration value. In both cases, at ε = 0 the number of components is equal to the number of nodes, n. As the filtration value increases, the number of components reduces, and eventually will reach a single one containing all nodes. Brain views generated with brain-net (Xia et al., 2013), r stands for Pearson’s correlation.

In this work, Topological Data Analysis is applied to explore individual brain networks based on the resting state functional MRI (rsfMRI) of children diagnosed with ADHD and typically developing children (TDC), obtained from the publicly available ADHD-200 database (Milham et al., 2012). First, the consistency of this methodology was explored when using four different brain segmentation schemes (atlases), and then group differences were identified between ADHD and TDC groups, at the whole-brain and sub-network levels. ADHD is a developmental disorder characterized by a lack of control of appropriate behavior and a difficulty to maintain attention (WHO, 1992; APA, 1994). Current theories propose the potential alteration of multiple functional networks and their interaction, including the default, cognitive control (fronto-parietal), dorsal and ventral attention, and salience networks (Sonuga-Barke and Castellanos, 2007; Castellanos and Aoki, 2016). Consequently, it was expected that the proposed methodology would reveal significant differences between groups among the components of these functional networks.

## 2. Methods

### 2.1 Sample

Imaging and phenotypic data from 263 participants corresponding to the New York University Child Study Center dataset were obtained from the ADHD-200 database (http://fcon1000.projects.nitrc.org/indi/adhd200/). Subjects reported with a secondary diagnosis and/or not medication-naïve status were discarded. Only those with good imaging quality and complete phenotypic information were used for subsequent analysis, resulting in a total of 182 children. Study protocols were approved by the New York University Institutional Review Boards, and after an explanation of study procedures a written informed consent from parents and assent from children were required.

Pediatric diagnosis was based on the Schedule of Affective Disorders and Schizophrenia for Children Present and Lifetime Version (KSADS-PL) and the Conners’ Parent Rating Scale-Revised, Long version (CPRS-LV). Moreover, IQ was measured with the Wechsler Abbreviated Scale of Intelligence (WASI). Inclusion in the ADHD group was based on the parent and child responses to KSADS-PL and obtaining a t-score greater or equal than 65 in any of the ADHD related indices of the CPRS-LV. TDC had ADHD summarized t-scores below 60, and lack of any DSM-IV axis-I disorders. Exclusion criteria were an IQ below 80 or any chronic medical conditions. However, phenotypic data of three ADHD datasets showed full intelligence scores below 80, while two TDC showed t-scores greater than 60 in the ADHD summary scale. Data from those subjects were discarded for further analysis, resulting in a final sample of 81 ADHD children (average age ±standard deviation: 10.5 ±2.48 years old) and 96 TDC (12.26 ±3.07 years old) (Table 1).

**Table 1.**
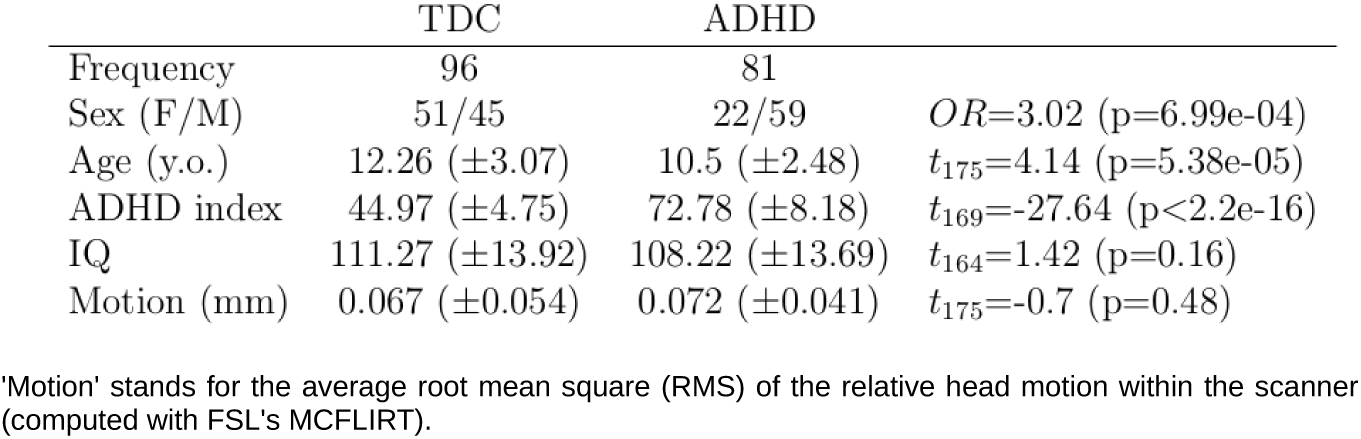
Phenotypic information by diagnostic group.

|  | TDC | ADHD |  |
| --- | --- | --- | --- |
| Frequency | 96 | 81 |  |
| Sex (F/M) | 51/45 | 22/59 | $OR=3.02$ ($p=6.99e-04$) |
| Age (y.o.) | 12.26 ($\pm 3.07$) | 10.5 ($\pm 2.48$) | $t_{175}=4.14$ ($p=5.38e-05$) |
| ADHD index | 44.97 ($\pm 4.75$) | 72.78 ($\pm 8.18$) | $t_{169}=-27.64$ ($p<2.2e-16$) |
| IQ | 111.27 ($\pm 13.92$) | 108.22 ($\pm 13.69$) | $t_{164}=1.42$ ($p=0.16$) |
| Motion (mm) | 0.067 ($\pm 0.054$) | 0.072 ($\pm 0.041$) | $t_{175}=-0.7$ ($p=0.48$) |

'Motion' stands for the average root mean square (RMS) of the relative head motion within the scanner (computed with FSL's MCFLIRT).

**Table 2.** Descriptive statistics for the minimum filtration value to reach a single component, for each group and atlas.

| | $\bar{x}(sd)$ | range | $t_{175}(p)$ | Cohen's $d$ |
| --- | --- | --- | --- | --- |
| AAL-TDC | 0.573 (0.056) | 0.44 - 0.702 | 0.662 (0.509) | 0.101 |
| AAL-ADHD | 0.568 (0.062) | 0.452 - 0.803 |  |  |
| CC200-TDC | 0.558 (0.073) | 0.382 - 1 | 1.52 (0.13) | 0.228 |
| CC200-ADHD | 0.543 (0.059) | 0.423 - 0.715 |  |  |
| P264-TDC | 0.585 (0.07) | 0.479 - 1 | 0.334 (0.739) | 0.051 |
| P264-ADHD | 0.581 (0.089) | 0.505 - 1 |  |  |
| CC400-TDC | 0.566 (0.089) | 0.458 - 1 | 2.273 (0.024) | 0.338 |
| CC400-ADHD | 0.541 (0.044) | 0.434 - 0.645 |  |  |

### 2.2 Imaging acquisition

Magnetic resonance images were acquired with a Siemens Magnetom Allegra 3T scanner (Siemens Medical Solutions, Erlangen, Germany). Whole brain fMRI volume images were obtained using a T2*-weighted echo planar imaging interleaved sequence (TR/TE = 2000/15ms, flip angle = 90, voxel size 3×3×4 mm^3^, FOV = 240×192 mm^2^) with a scan duration of 6 minutes. Participants were instructed to remain still, close their eyes, think of nothing systematically and not fall asleep. In order to obtain an anatomical reference, high resolution structural T1-weighted Magnetization Prepared Rapid Acquisition Gradient Echo (MPRAGE) images were acquired (TR/TE = 2530/3.25 ms, flip angle = 7^°^, voxel size 1.3 × 1.0 × 1.3 mm^3^, FOV = 256 × 256 mm^2^).

### 2.3 Preprocessing

Preprocessing was implemented using FMRIB’s Software Libraries (FSL v.5.0.6) (Jenkinson et al., 2012). Steps included removing the first four volumes, slice timing, head motion correction, brain extraction, regression of confounding variables, band-pass temporal filtering (0.01–0.08 Hz), and spatial normalization. Given that psychiatric and pediatric populations usually show higher in scanner motion than controls and adults (Satterthwaite et al., 2012), a rigorous confounding regression strategy was implemented to minimize head motion artifact. Specifically, several variables were regressed out from the functional data, including the six rigid-body motion parameters, the average signal from both white matter (WM) and cerebrospinal fluid (CSF), the derivative of these eight parameters and the square of these sixteen variables (Gracia-Tabuenca et al., 2018). In addition, to minimize the impact of physiological noise, five principal components of the signal from WM and CSF were also included as confounding variables (Behzadi et al., 2007; Chai et al., 2012). Furthermore, those volumes with a root mean square (RMS) of relative head motion greater than 0.25 mm were also included as confounds (Satterthwaite et al., 2013). Subjects with an average RMS of relative head motion higher than 0.55 mm or less than 4 minutes of non-motion-affected data, were discarded. Eventually, each fMRI volume was registered to its corresponding T1 image with a rigid-body transformation, followed by an affine and non-lineal registration to a 2 × 2 x 2 mm^3^ children-specific template, the 4.5–18.5 years old NIHPD atlas (Fonov et al., 2011).

### 2.4 Functional connectomes

For every dataset, four functional connectomes (connectivity matrices) were computed based on different brain atlases: AAL (Tzourio-Mazoyer et al., 2002), P264 (Power et al., 2011), CC200, and CC400 (Craddock et al., 2012). All of them include cerebrum and cerebellum. The first one consists of a segmentation of 116 anatomical regions, the second one is a set of 264 spherical ROIs with high reliability in both task and resting fMRI large datasets, while the last two are segmented based on functional connectivity homogeneity (with 190 and 351 nodes, respectively).

For each subject and atlas, the average fMRI signal of every defined region was extracted and then the functional connectome was computed as the Pearson’s cross-correlation between all possible pairs of regions. The variability of the explored TDA variables along the atlases was assessed by the Kendall’s Concordance Coefficient (KCC).

### 2.5 Topological Data Analysis (TDA)

Given a set of nodes V and a measure x_i_ corresponding to the i-th node, we define the set F = {x_1,_ x_2,_ …, x_n_}. For a positive number ε we state that two nodes in F are connected if their distance is less than ε, the filtration value. A k-simplex is a subset of k-1 nodes in F pairwise connected, then a node is a 0-simplex, a 1-simplex is an edge, a 2-simplex is a triangle, and so on. The corresponding Rips complex, denoted by Rips(F,ε), is a collection of k-simplices obtained for the filtration value ε. For every sequence of filtration values ε_1_, ε_2_, …, ε_j_ there is a nested sequence of simplicial complexes Rips(F,ε_1_) ⊂ Rips(F,ε_2_) ⊂ … ⊂ Rips(F,ε_j_) which is called Rips filtration. Algebraic information extracted from this topological space are called Betti numbers, particularly, the Betti-0 number (B_0_) accounts for the number of components, i.e. the number of isolated nodes or sets of nodes connected by a sequence of edges; Betti-1 number refers to the number of cavities in the 2-dimensional space between nodes, and so on (for more extensive reviews on topological data analysis see Edelsbrunner, 2000; and Sizemore et al., 2019). In this work, we focus exclusively on B_0_. If we start with a filtration value ε=0, all nodes are disconnected, and the number of components is the number of 0-simplices (nodes). When ε increases, some isolated nodes will connect with others and the number of components decreases. Therefore, B_0_ will diminish as the nodes gradually connect to each other as ε increases. It is possible to identify the filtration values for which there is a change in B_0_, until there is only one large component containing all the nodes. This process is summarized in the so called B_0_ curve (Figure 1).

Here, the distance between nodes is defined as in Lee et al (2012), i.e., d(x_i_, x_j_) = 1-r(x_i_, x_j_), where r is the Pearson’s correlation between the pair of nodes x_i_ and x_j_. The B_0_ curves are computed with the TDA package in R (https://cran.r-project.org/web/packages/TDA/index.html), and characterized in terms of the area under the curve, slope and kurtosis. The area under the curve accounts for the overall transition from all nodes being isolated to being connected into a single component, with smaller areas suggesting that B_0_ decreases with smaller filtration values. The slope accounts for the rate of change, being all negative, lower values mean a faster transition to a single component. Finally, the kurtosis accounts for how “tailed” the distribution is with respect to the average value, with higher values meaning faster transition to a single component.

### 2.6 Group inferences

Differences between ADHD and TDC groups were assessed for the filtration value when the brain network reached the single component containing all nodes. In addition, the explored properties of the B_0_ curves were tested with a logistic regression including sex, age and average RMS of the relative head motion as confounding variables. All dimensional variables were standardized to z-scores. Both approaches were tested for each of the four brain atlases.

Logistic regression was also assessed for every intra- and inter-network combination of nodes. This approach was independently implemented for the seven lobes of the AAL atlas, and the thirteen functional networks defined in the P264 atlas. Given the 28 and 91 possible combinations, respectively, significance was set to p < 0.05, corrected for a false discovery rate (FDR), q < 0.05 (Benjamini and Hochberg, 1995).

## 3. Results

### 3.1 Agreement across brain atlases

The three explored properties of the B_0_ curves showed a generalized sample agreement along the four brain parcellations. Significant agreement was found considering every atlas and each feature: area (KCC = 0.87; X^2^_176_ = 609; p = 4.67e-49), slope (KCC = 0.68; X^2^_176_ = 477; p = 1.33e-29) and kurtosis (KCC = 0.44; X^2^_176_ = 307; p = 3.65e-09). Moreover, pairwise concordance coefficient was significant for every pair of atlases and every TDA metric (Figure 2; Supplementary Tables 1-3).

**Figure 2.**
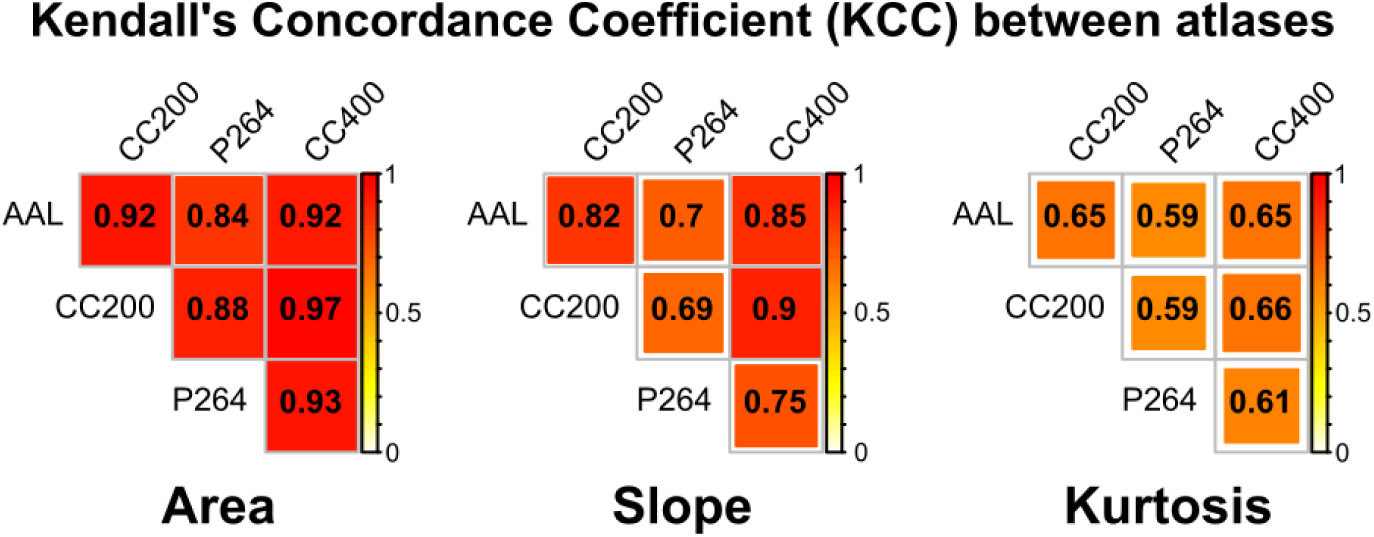
Kendall’s Concordance Coefficient (KCC) between brain parcellations for the explored properties of the B_0_ curves: area under the curve, slope and kurtosis. KCC value is depicted in yellow-red, with p(KCC > 0.59) < 0.05, given two raters and 176 degrees of freedom.

### 3.2 Whole brain topology

The average minimum filtration value for which the large single component was connected was in the range between 0.54 and 0.59 for all atlases and groups. However, for every atlas, the mean filtration value was higher for the TDC group, being statistically significant for the CC400 atlas (t_175_ = 2.273; p = 0.024; d = 0.338). This means that the ADHD functional connectomes tend to reach the single component with lower filtration values compared to those of the TDC.

The area under the B_0_ curves showed significantly lower odds for the ADHD group, no matter the brain atlas (0.572 < OR < 0.622; 0.0028 < p < 0.014; Supplementary Table 4), which means that the ADHD group has smaller areas compared to the TDC group (Figure 3). The area under the B_0_ curves accounts for the overall transition from all nodes being isolated to being connected into a single component, with smaller areas when B_0_ decreases faster as the filtration value increases. In other words, less area under the curve implies lower number of components, i.e. less segregation, which should be mediated by increased connectivity in the edges mediating the integration of components. In order to explore such edges, the proportion of subjects showing connectivity was compared between groups for each edge at some filtration values (Figure 4). These tests showed widespread frontal short-range and cortical long-range edges being more frequently present in the ADHD group (p<0.01, uncorrected; Figure 4). No group differences were found for the slope nor the kurtosis (Supplementary Table 4).

**Figure 3.**
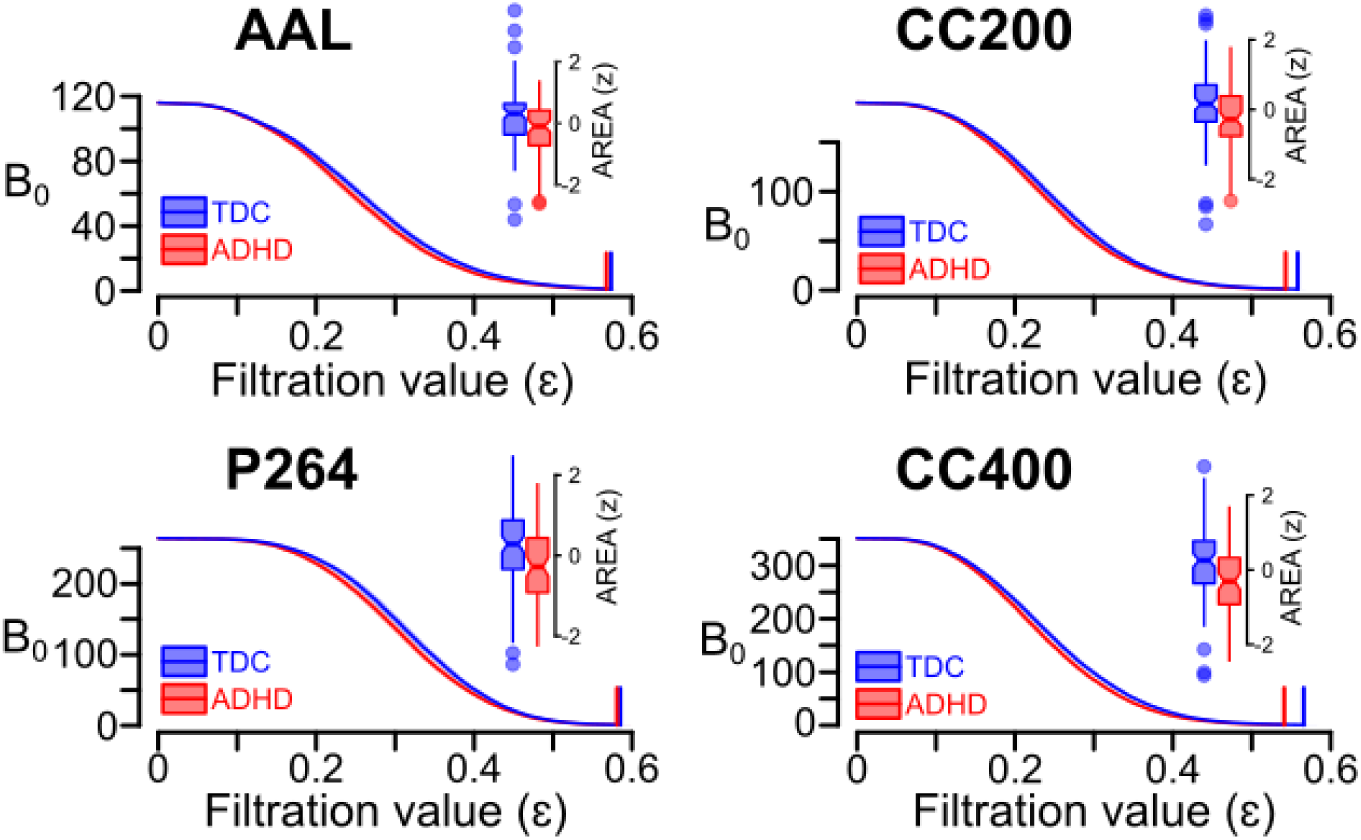
B_0_ curves for each group and brain parcellation. Group average with 95% confidence interval of the B_0_ curves. Notched boxplots of the area under the curve (z-values) are depicted for each brain atlas: AAL (top-left), CC200 (top-right), P264 (bottom-left), and CC400 (bottom-right). Vertical lines on the B_0_ curves depict the average filtration value at which all the nodes connect into a single component.

**Figure 4.**
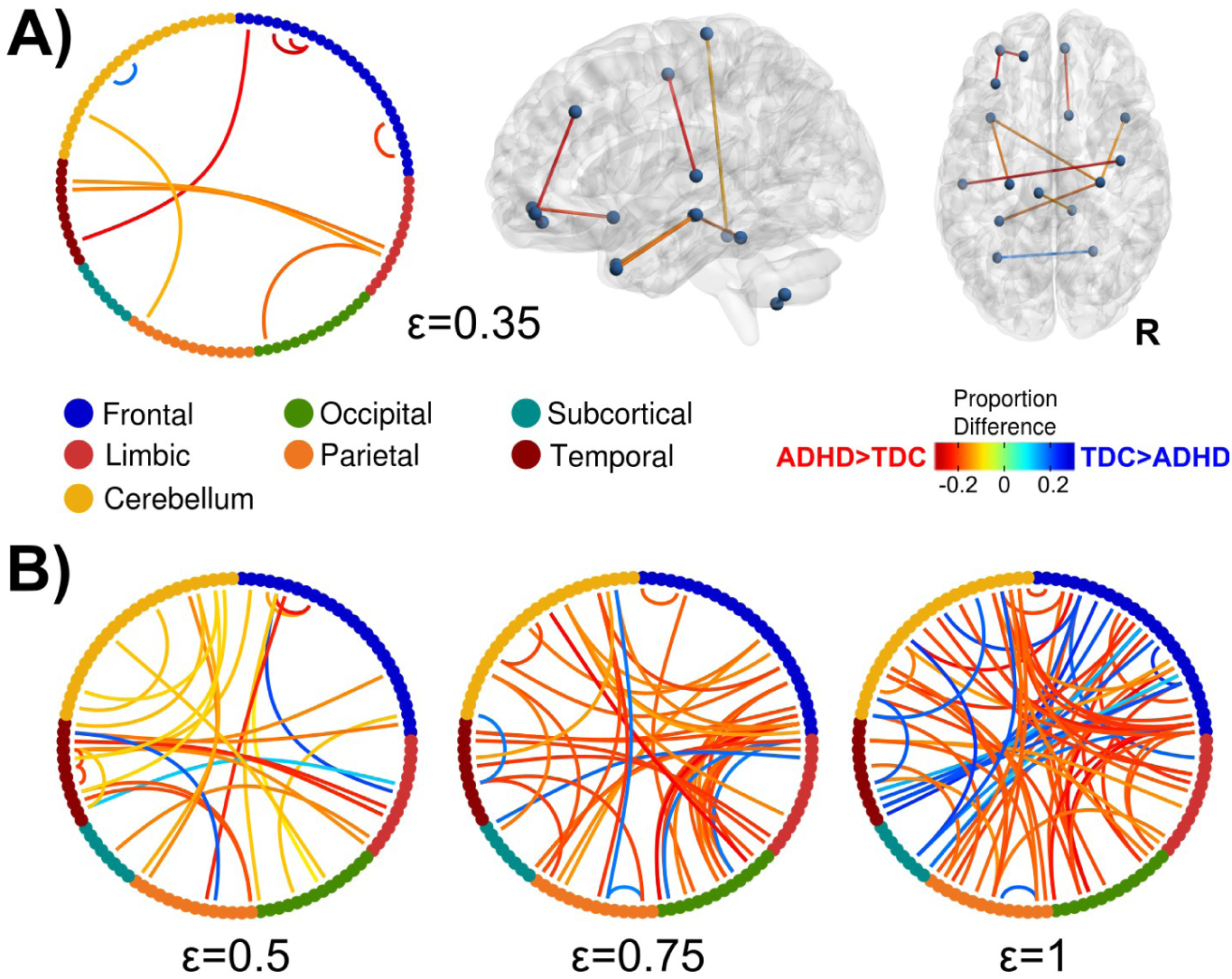
Edges with differences in the proportion of subjects between groups at four different filtration values (ε = 0.35, 0.5, 0.75, and 1). Nodes from each lobe (AAL atlas) are represented with different colors in the chord diagrams. Only edges with a proportion difference at p < 0.01 (uncorrected) are depicted. For ε = 0.35 the edges are represented in the brain using brain-net (Xia et al., 2013). R stands for the right side of the brain.

### 3.3 Intra- and inter-network inference

The same analyses were performed in subsets of nodes, corresponding to the nodes of a single lobule or functional network (intra-network) or the nodes of two networks (inter-network), according to the lobular and the functional parcellation of the AAL and P264 atlases, respectively. Only those combinations including the frontal lobe showed significant differences for the area under the B_0_ curves (FDR corrected at q < 0.05).

These subsets included the intra-network and every possible inter-network subset including the frontal lobe, and in every case the ADHD group showed a lower area compared to the TDC (Table 3; Figure 5). A subset of nodes including the temporal and subcortical nodes showed marginally significant differences (p < 0.05, uncorrected), also evidence for less area for the ADHD group. These results demonstrate a widespread decreased segregation of the brain network in the ADHD group, particularly involving the frontal lobe. When considering the functional systems in the P264 atlas, notably, all the subsets of nodes that included the DMN also showed smaller areas for the ADHD group, but other intra- and inter-network subsets showed similar patterns (Figure 5). However, these results were marginally significant (p < 0.05, uncorrected) given the higher number of tests to correct for (n=91).

**Table 3.** Logistic regression odds ratios (OR) for the area under the B_0_ curves for the subsets of AAL nodes with significant differences between groups (ADHD < TDC)

| | OR | $\log(\text{OR})$ | p |
| --- | --- | --- | --- |
| Frontal-Frontal | 0.477 | -0.741 | 6.32e-04 |
| Frontal-Limbic | 0.571 | -0.56 | 0.0067 |
| Frontal-Occipital | 0.604 | -0.504 | 0.01 |
| Frontal-Parietal | 0.473 | -0.748 | 4.52e-04 |
| Frontal-Subcortical | 0.472 | -0.75 | 5.00e-04 |
| Frontal-Temporal | 0.526 | -0.642 | 0.0025 |
| Frontal-Cerebellum | 0.582 | -0.542 | 0.0051 |

$p < 0.05$ , FDR-corrected.

**Figure 5.**
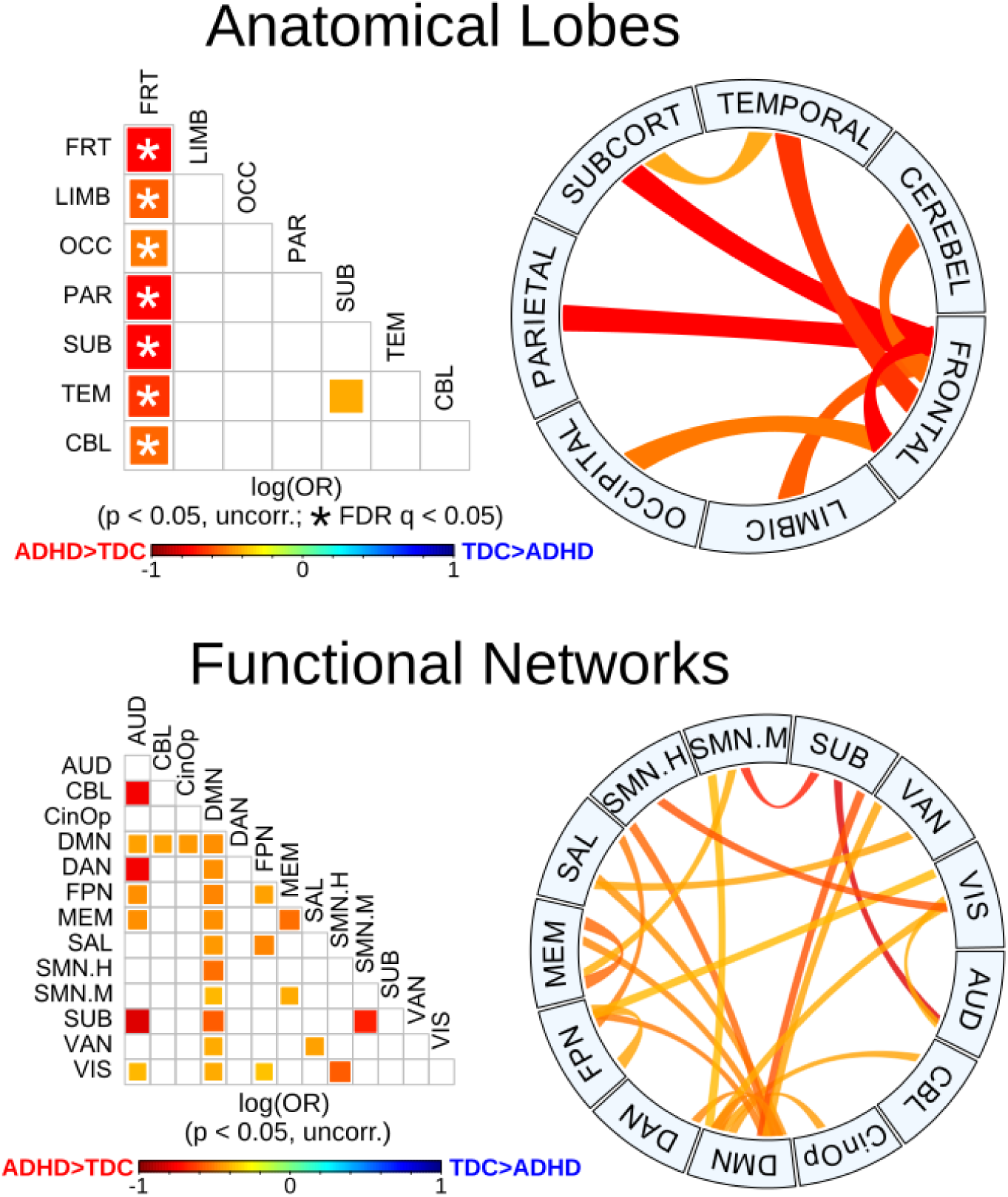
Group differences for anatomical and functional sub-networks. Pairwise plot and chord diagrams (Gu et al., 2014) of significant differences for the area under the B_0_ curves (p < 0.05) between groups. Anatomical lobes (top) are based on AAL parcellation and functional networks (bottom) are based on P264 parcellation. * denotes FDR-corrected (q < 0.05) in the pairwise plots. Abbreviations: auditory (AUD), cerebellar (CBL), cingulo-opercular (CinOp), default mode (DMN), dorsal attention (DAN), fronto-parietal (FPN), memory retrieval (MEM), salience (SAL), sensory/somatomotor hand (SMN.H), sensory/somatomotor mouth (SMN.H), subcortical (SUB), ventral attention (VAN), visual (VIS).

## 4. Discussion

In this work, methods from Topological Data Analysis were applied to explore the topology of the brain network as a function of the filtration value (i.e., the connectivity threshold). Resulting B_0_ curves were characterized in terms of three parameters, the area under the curve, slope and kurtosis, and compared between ADHD and TDC. The consistency across four brain segmentation schemes was highest for the area, then the slope and lowest for the kurtosis. In addition, application of this model to a pediatric sample showed that the area under the curve was significantly lower for the ADHD group, both at the whole brain and at the sub-network levels. These results showed decreased functional segregation in the ADHD group, mainly involving the frontal lobe and the default mode network.

Pairwise agreement between brain parcellations was high for the area under the curve (KCC range: 0.83-0.97) and the slope (KCC range: 0.68-0.9), and medium to low for the kurtosis (KCC range: 0.57-0.67). These results suggest that this methodology is consistent among different parcellation schemes, especially for the area under the B_0_ curve. In addition, considering that the Rips filtration does not depend on a particular connectivity threshold, but instead explores all the filtration values with a change in the topology of the network, this methodology contributes to provide a more complete picture of the brain network, overcoming one of the main limitations of other approaches. Taken together, these are potentially important advantages compared to other methods applied to brain networks, such as graph theory, which has been shown to be highly dependent on the brain parcellation scheme (Wang et al., 2009a; Chen et al., 2018; Doucet et al., 2019), and on the selection of a connectivity threshold or connectivity cost (van den Heuvel et al., 2008; Fornito et al., 2010; Tomasi and Volkow, 2010; Gracia-Tabuenca et al., 2018; Termenon et al., 2016).

The area under the B_0_ curve was significantly lower for the ADHD group, both at the whole brain network and at the sub-network level, being strikingly significant for the interactions involving the frontal lobe. As mentioned above, the area under the curve accounts for the overall transition from all nodes being isolated to being connected into a single component, with smaller areas suggesting that B_0_, the number of components, decreases faster as the filtration value increases. Such differences in B_0_ for a given filtration value are mediated by edges that bind together previously split components, which results from increased connectivity in some edges mediating the integration into larger components. Taken together, the results here presented can be interpreted as higher functional connectivity within the brain network and specific sub-networks in the ADHD group, especially those involving the frontal lobe. Previous evidence has also suggested increased functional connectivity in a variety of regions of the frontal lobe in ADHD (Tomasi and. Volkow, 2012; Hoekzema et al., 2014; Mostert et al., 2016), as well as fronto-occipital (Cocchi et al., 2012) and fronto-subcortical connections (Cocchi et al., 2012; Tomasi et al., 2012), particularly those associated with reward and motivation (Tomasi and Volkow, 2012). In addition, our results showed a similarly widespread pattern in several functional sub-networks, mainly the default mode, but also attention, salience, fronto-parietal and auditory nodes, among others (Figure 5). Although the latter results did not survive FDR correction, they provide the basis to infer the potential functional systems being affected in ADHD, being consistent with the current theories involving such networks (Castellanos and Aoki, 2016), particularly with the DMN interference hypothesis, which is based on the findings of altered interactions between the DMN and networks involved in top-down executive control (Fox et al., 2005, 2007; Kelly et al., 2008; Castellanos and Aoki, 2016; Elton et al., 2014; Hoekzema et al., 2014; Bos et al., 2017; Qian et al., 2019).

Previous studies have reported a myriad of differences in network properties between ADHD and TDC participants. At the whole-brain level, higher functional segregation and lower functional integration in ADHD subjects compared to controls have been reported (Wang et al., 2009b; Lin et al., 2014), although other groups did not reproduce those results (Cocchi et al., 2012; Sato et al., 2013). Since the decreased area under the B_0_ curves could be interpreted as higher integration and lower segregation of isolated components, our results seem to be contradictory to the aforementioned ones. Nevertheless, the previous studies explored connectivity costs higher than 10%, which according to Lin et al., (2014) would correspond to filtration values higher than ε = 0.5, when most of the subjects actually exhibit a single component (Figure 3). Therefore, these results are actually complementary, given that B_0_ curves consider a wider range of connectivity thresholds, rarely explored with graph theory. Indeed, when exploring the edge-wise proportions between groups at different connectivity thresholds (Figure 4), the ADHD group showed consistently widespread increases compared to the TDC. However, at lower connectivity thresholds (higher filtration values), the ADHD group showed decreased proportion of edges in several interactions, mainly including the frontal, temporal, subcortical and cerebellar regions, which seem consistent with previous reports of decreased connectivity in ADHD (Tomasi and Volkow, 2012; Di Martino et al., 2013; Elton et al., 2014). These results evidence that the static representation of the network changes as a function of the connectivity threshold, therefore an approach that takes into account wider threshold ranges should provide better insights into the neurophysiological substrate of ADHD.

As far as we are concerned, only two previous studies have explored B_0_ in ADHD brain networks (Lee et al., 2011, 2012), using fludeoxyglucose positron emission tomography (FDG-PET) and inter-region covariation at the sample level, qualitatively reporting higher number of components for the ADHD group compared to the TDC. Such findings seem to be opposite to the results here presented; however, methodological differences prevent direct comparisons between results. First, time-scales are significantly different, with the FDG-PET scans reflecting the glucose uptake occurring during several minutes; in contrast, rsfMRI reflects variations in blood oxygenation during tens of seconds. Furthermore, FDG-PET connectivity matrices reflect inter-region covariation of (long term) glucose metabolism across subjects, while rsfMRI connectivity matrices reflect inter-region covariation (within seconds) within the same subject and later compared between groups. Overall, both methodologies potentially reflect complementary aspects of the functional connectomes in ADHD.

## Conclusion

In summary, the present study showed a robust and informative implementation of topological data analysis in functional connectomics. The results exhibited significant differences for the brain topology of children with ADHD, both at the whole brain network and at the functional sub-network levels, particularly involving the frontal lobe and the DMN. Therefore, this approach may contribute to identifying the physio-pathology of neurodevelopmental disorders, reducing the bias of connectomics-related methods selection.

## Acknowledgments

We are extremely grateful to the ADHD-200 consortium for such an important data sharing initiative. We also thank Leopoldo Gonzalez-Santos for his technical support and Michael C. Jeziorski for editing the manuscript. Zeus Gracia Tabuenca is a doctoral student at the “Programa de Doctorado en Ciencias Biomédicas, Universidad Nacional Autonoma de Mexico (UNAM)” and received a fellowship (330142) from “Consejo Nacional de Ciencia y Tecnología” (CONACYT). CONACYT had no role in study design, data collection, analyses nor writing the manuscript.

