## Supplementary Table 1 for "Topological Data Analysis reveals robust alterations in the whole-brain and frontal lobe functional connectomes in Attention-Deficit/Hyperactivity Disorder"

Supplementary Table 1. Pairwise Kendall's coefficient of concordance (KCC) between atlases for the area under the filtration curve.

| | KCC | $\chi^2_{176}$ | p |
|---|---|---|---|
| AAL-CC200 | 0.92 | 324 | 7.52e-11 |
| AAL-P264 | 0.838 | 295 | 4.97e-08 |
| AAL-CC400 | 0.916 | 323 | 1.05e-10 |
| CC200-P264 | 0.884 | 311 | 1.46e-09 |
| CC200-CC400 | 0.973 | 342 | 8.56e-13 |
| P264-CC400 | 0.928 | 327 | 3.9e-11 |
