## Supplementary Table 2 for "Topological Data Analysis reveals robust alterations in the whole-brain and frontal lobe functional connectomes in Attention-Deficit/Hyperactivity Disorder"

Supplementary Table 2. Pairwise Kendall's coefficient of concordance (KCC) between atlases for the kurtosis of the filtration curve.

| | KCC | $\chi^2_{176}$ | p |
|---|---|---|---|
| AAL-CC200 | 0.648 | 228 | 0.0051 |
| AAL-P264 | 0.591 | 208 | 0.049 |
| AAL-CC400 | 0.648 | 228 | 0.0051 |
| CC200-P264 | 0.593 | 209 | 0.046 |
| CC200-CC400 | 0.655 | 231 | 0.0035 |
| P264-CC400 | 0.609 | 214 | 0.026 |
