## Supplementary Table 3 for "Topological Data Analysis reveals robust alterations in the whole-brain and frontal lobe functional connectomes in Attention-Deficit/Hyperactivity Disorder"

Supplementary Table 3. Pairwise Kendall's coefficient of concordance (KCC) between atlases for the slope of the filtration curve.

| | KCC | $\chi^2_{176}$ | p |
|---|---|---|---|
| AAL-CC200 | 0.821 | 289 | 1.7e-07 |
| AAL-P264 | 0.705 | 248 | 2.83e-04 |
| AAL-CC400 | 0.848 | 298 | 2.33e-08 |
| CC200-P264 | 0.69 | 243 | 6.13e-04 |
| CC200-CC400 | 0.9 | 317 | 4.03e-10 |
| P264-CC400 | 0.747 | 263 | 2.3e-05 |
