## Supplementary Table 4 for "Topological Data Analysis reveals robust alterations in the whole-brain and frontal lobe functional connectomes in Attention-Deficit/Hyperactivity Disorder"

Supplementary Table 4. Logistic regression odds ratio for every coefficient for each brain atlas, testing for differences between ADHD and TDC groups. 'Motion': average RMS head motion.

| | Area | Kurtosis | Slope | Sex | Age | Motion |
|---|---|---|---|---|---|---|
| AAL | 0.622 | 0.899 | 1.08 | 3.22 | 0.492 | 0.882 |
| (p-value) | (0.0141) | (0.564) | (0.677) | (8.83e-4) | (1.77e-4) | (0.882) |
| CC200 | 0.612 | 0.889 | 0.893 | 3.4 | 0.487 | 0.95 |
| (p-value) | (0.008) | (0.561) | (0.554) | (5.42e-4) | (1.38e-4) | (0.764) |
| P264 | 0.611 | 1.05 | 0.899 | 3.49 | 0.495 | 0.877 |
| (p-value) | (0.013) | (0.763) | (0.579) | (4.96e-4) | (1.45e-4) | (0.495) |
| CC400 | 0.572 | 0.719 | 1.13 | 3.33 | 0.521 | 0.871 |
| (p-value) | (0.003) | (0.351) | (0.517) | (7.79e-4) | (5.57e-4) | (0.431) |
